## Supplementary figures and images for "The spread of pESI-mediated extended-spectrum cephalosporin resistance in *Salmonella* serovars - Infantis, Senftenberg, and Alachua isolated from food animal sources in the United States"

### S1 Fig

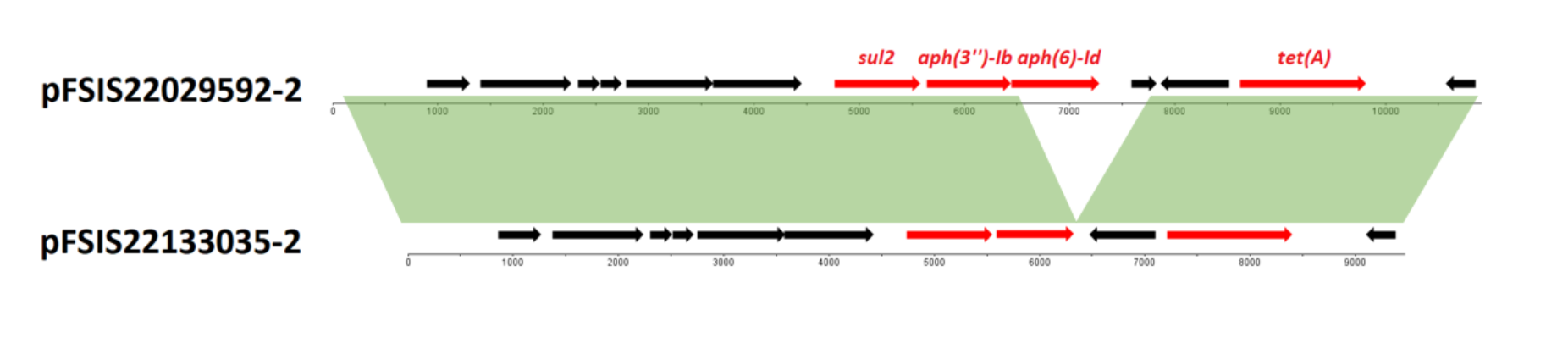

### S2 Fig

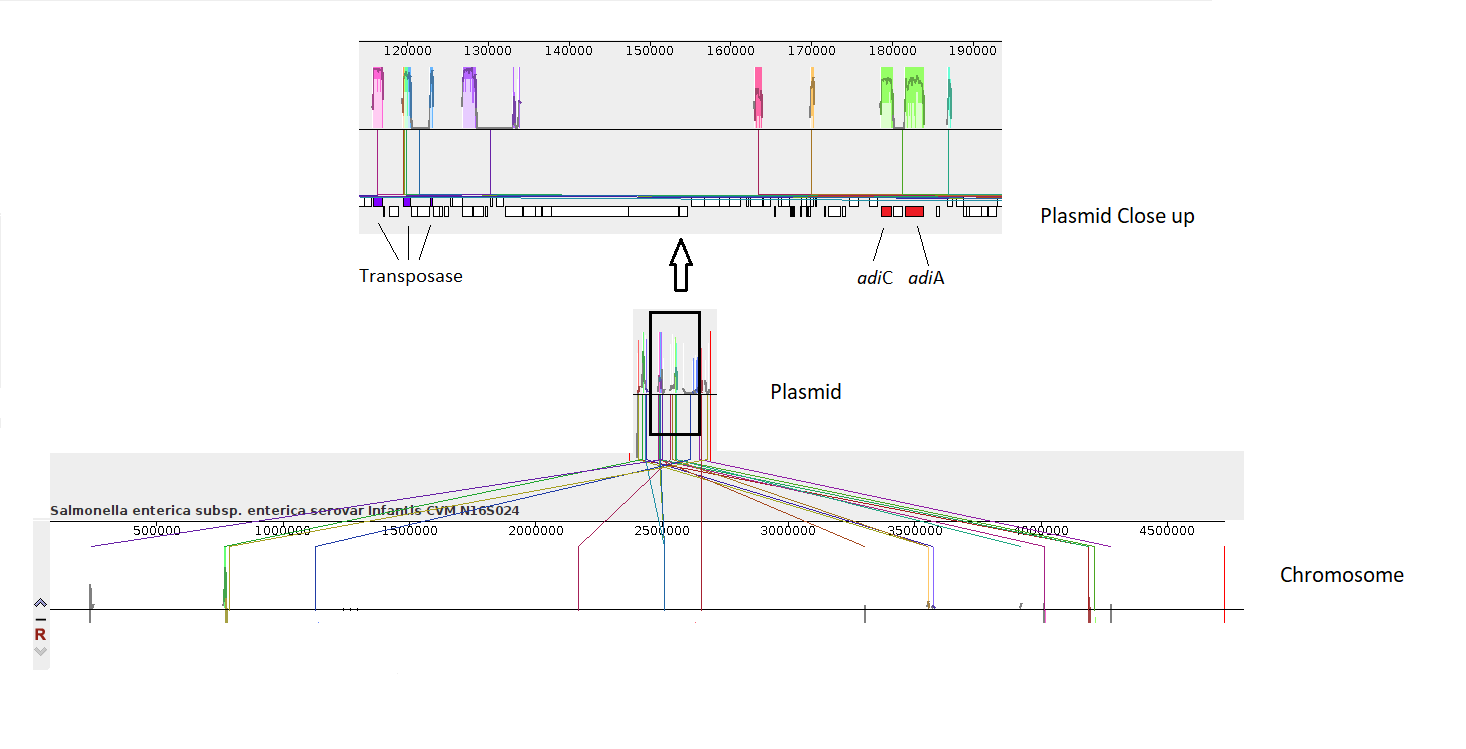

### S3 Fig

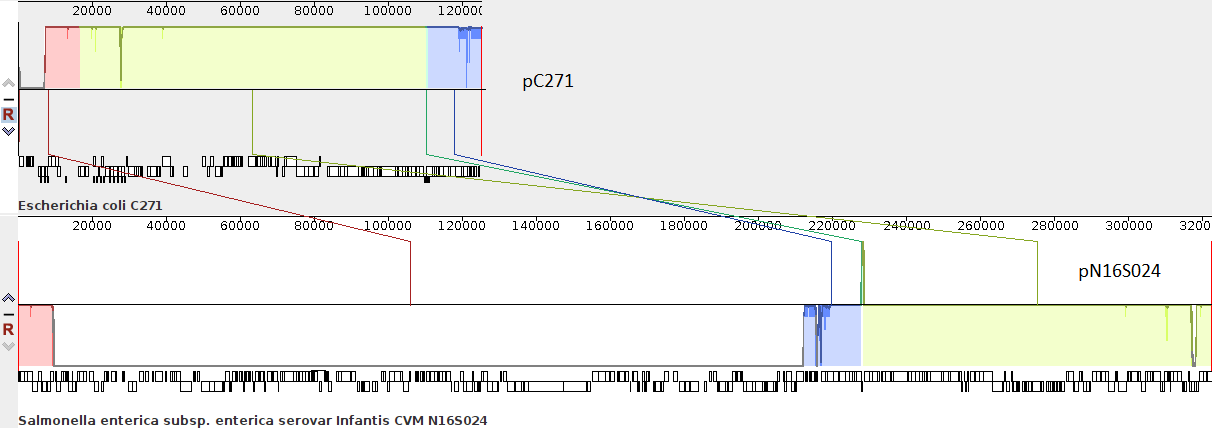
